# Effects of treatment with three antibiotics, vancomycin, neomycin, and AVNM on gut microbiome in C57BL/6 mice

**DOI:** 10.1101/2021.02.08.430372

**Authors:** Pratikshya Ray, Subhayan Chakraborty, Arindam Ghosh, Palok Aich

## Abstract

Higher organisms, especially mammals, harbor diverse microbiota in the gut that plays a major role in maintaining health and physiological homeostasis. Perturbation of gut flora helps identifying their roles. Antibiotics are potent perturbing agents of microbiome. Select antibiotics like vancomycin, neomycin, and AVNM (an antibiotic cocktail containing ampicillin, vancomycin, neomycin, and metronidazole) were used to perturb the gut microbiota of C57BL/6 male mice to understand their roles in host immunity and metabolism. The current study revealed that the resulting gut microbial composition was different, and diversity (at the phylum and genus level) was reduced differentially following each antibiotic treatment. Vancomycin treatment caused a significant increase in Verrucomicrobia and Proteobacteria phyla. The treatment with neomycin yielded an increase in the Bacteroidetes phylum, while the treatment with AVNM led to an increase in Proteobacteria phylum with lowest diversity of microbiome in the gut. The current results also revealed that the different antibiotics treatment caused variation in the cecal index, expression of immune genes (TNF-α, IL-10, IFN-γ) in the colon, and short-chain fatty acids (SCFA) level in the blood of mice. A strong correlation was observed for antibiotic-induced differential dysbiosis patterns of gut microbiota and the altered immune and SCFA profile of the host. The outcome of the present study could be clinically important.

## Introduction

The mammalian gastrointestinal tract is inhabited by a hundred trillions of highly diverse and complex microbes (1, 2). It was well established that the abundance and diversity of these large numbers of gut microbiota play an important role in regulating the immune response of the host (3, 4). To understand the role of specific gut microbes in the host, perturbation is the most effective way. Antibiotics are widely used as the most potent perturbing agent of gut microbiota (5–7). Each antibiotic altered the abundance of some specific groups of gut microbes (8).

Gut microbes produce metabolites like acetate, propionate, and butyrate that belong to the group of SCFAs by metabolizing various dietary fibers in the host (9). SCFAs could suppress the lipopolysaccharide (LPS) and pro-inflammatory cytokines like TNFα, IL-6, and might increase the production of anti-inflammatory cytokines, IL-10 (10, 11). During the dysbiosis of gut microbiota, the increase in the Proteobacteria group of bacteria may cause an increase in the blood endotoxin level through their LPS. LPS could enhance the production of various pro-inflammatory cytokines by activating different Toll-Like Receptors (TLR4) of the gut epithelial cells (12–16). It was observed that in the colon tissue, butyrate, an important SCFA, caused inhibition of the LPS induced activation of NF-κB (17–19). Dysbiosis of gut microbiota can be introduced by antibiotic treatment.

Dysbiosis, due to antibiotic treatment, could reduce the diversity and may change the composition of the gut microbiota to lead to a pathogenesis like inflammatory bowel disease (IBD) (20). A significant increase in certain opportunistic pathogens, resistant bacteria like *Clostridium difficile,* and a decrease in beneficial butyrate-producing bacteria were observed in IBD patients compared to healthy individuals (21). The samples from fecal material of IBD patients and long term antibiotic using individuals contained very less amount of SCFAs that could lead to a higher range of inflammation in the host (5, 9, 22)<u>. The</u> restoration of gut microbiota following cessation of long-term antibiotic therapy was a time taking incomplete process (23). The correlation between the antibiotic-induced alterations of specific groups of microbes with the immune and metabolic response of the host is still not clear in the literature. Therefore, treatment with single and different combinations of select antibiotics can give us ideas about the extent of perturbation of specific group of gut bacteria and their effects on the host immune and metabolic response. Recently, we reported that treatment with vancomycin in C57BL/6 and BALB/c mice could induce dysbiosis till day 4 following treatment but continued treatment with the antibiotic confers physiological benefit by increasing *A. muciniphila* of verrucomicrobia phylum (24, 25).

In the current study, we compared the efficiency of perturbation of mouse gut microbiota either by vancomycin or neomycin or a cocktail known as AVNM containing ampicillin, vancomycin, neomycin, and metronidazole. Vancomycin is a broad-spectrum antibiotic to treat MRSA or drug-resistant *clostridium difficile* induced colitis (26, 27). Neomycin is an aminoglycosidic antibiotic that arrests the growth of intestinal bacteria (28). Neomycin is an antibiotic that is very effective against gram negative bacteria of proteobacteria phylum. By killing bacteria in the gut, it keeps the ammonia level low to prevent hepatic encephalopathy. It works well against streptomycin-resistance bacteria. While neomycin cannot be given intra-venous for its renal toxicity but vancomycin is given intra-venous (29–32). AVNM, the mixture of four different antibiotics, is well established as a gut microbiota depleting agent in mice (33, 34). The role of gnotobiotic or germ-free mice models are well documented and important model system in microbiome study, but because of the non-availability or inaccessibility of this model to a wide variety of scientific community, the AVNM treated mouse model serves as a good alternative.

Because of the different structures and functions of the antibiotics used, the treatment with each type of antibiotic caused a different kind of gut microbial modulation. Moreover, the alteration of gut microbes due to different antibiotics treatment caused the differential immune and metabolic response in the host. Expression of various pro- and anti-inflammatory immune genes and the production of specific SCFAs were strongly correlated with the abundance of specific phyla of gut microbes. The cecal size of mice also got differentially affected following treatment with different antibiotics.

## Results

### Antibiotic treatment alters the abundance and diversity of gut microbiota

Earlier reports showed the differential effects of treatment with several antibiotics causing dysbiosis of gut microbiota (5). A comparative analysis of select antibiotics to understand the changes in the composition of gut microbiota is warranted to correlate with innate mucosal immunity and systemic metabolites. The current results revealed that the gut microbiota of untreated C57BL/6 mice majorly contained Firmicutes and Bacteroidetes phyla with a very low percentage of Proteobacteria phylum (Fig 1A). Following treatment with vancomycin for seven consecutive days caused an increase in Verrucomicobia (by 71%) and Proteobacteria (by 20%) with a concomitant decrease in Firmicutes and Bacteroidetes phyla (Table 1). On the contrary, treatment with neomycin for seven days caused a significant increase in Bacteroidetes (by 72%) and a decrease in Firmicutes like major phylum (by 23%) (Table 1). Treatment with AVNM, in accordance, caused a significant increase in Proteobacteria (by 80%) and a decrease in major phyla like Firmicutes and Bacteroidetes (Fig 1A). Genus level analysis further validated the phylum level observation.

**Table 1:** Percent abundance of major phyla of gut microbes in the untreated control and different antibiotics treated mice.

|  | % Abundance |  |  |  |
| --- | --- | --- | --- | --- |
|  | Firmicutes | Bacteroidetes | Proteobacteria | Verrucomicrobia |
| Control | 52(±5) | 38(±4) | 1.8(±0.3) | 1.2(±0.4) |
| Vancomycin | 9(±2) | NIL | 20(±4) | 71(±6) |
| Neomycin | 23(±4) | 72(±5) | 1(±0.5) | NIL |
| AVNM | NIL | 2(±0.9) | 80(±4) | 18(±3) |

Major groups of bacterial phyla present in the cecal content of mice are determined by metagenomic analysis (16s rRNA) at different conditions (vancomycin, neomycin, AVNM treated groups along with the time-matched control) of C57BL/6 mice. Means of percent abundance of various phyla with their respective standard deviations (±SD) are shown.

**Fig 1.**
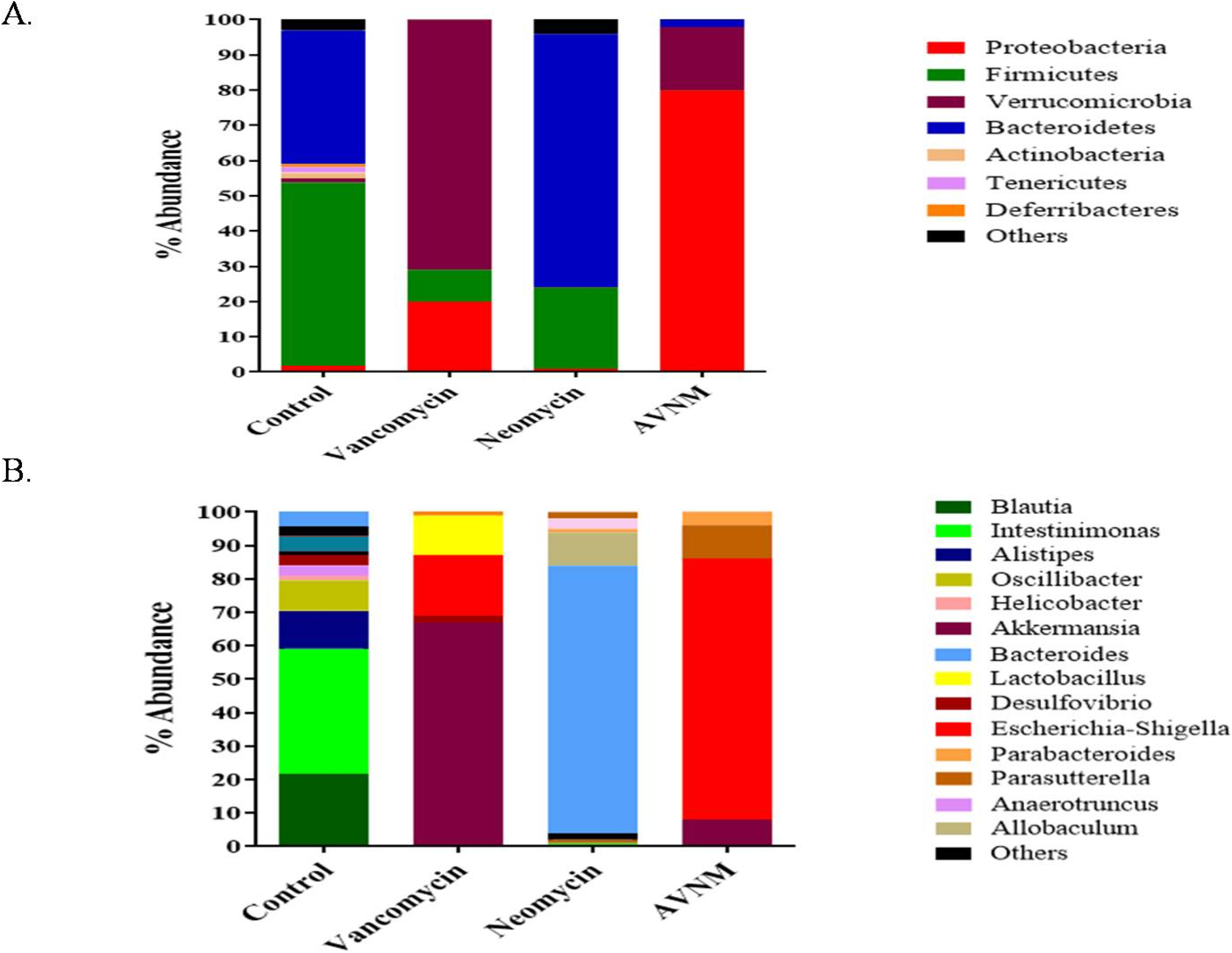
Composition of gut microbiota at phylum and genus level for control and antibiotics treated mice. Percent abundance of the gut microbiota composition in untreated (control) and antibiotics (vancomycin, neomycin, AVNM) treated mice following 7 days of treatment for A. phylum and B. genus level are shown. The percent abundance is calculated from the average values of at least 3 replicates. To avoid clutter the standard deviation (which is within 10%) of the data is not shown.

Genus level data showed that vancomycin treatment caused mostly an increase in the Akkermansia genus of Verrucomicrobia phylum (Fig 1B). While neomycin treatment caused an increase in the Bacteroides genus of Bacteroidetes phylum. However, AVNM treatment showed mostly elevation of Escherichia-Shigella genera of Proteobacteria phylum in the gut (Fig 1B).

Other members of the phylum or genus shown in the figures are for depicting the overall idea of composition and qualitative diversity of the gut microbiota. A detailed analysis of diversity is described below.

### Alpha diversity of gut microbiota decreased following treatment with antibiotics

Measurement of diversity is one of the important parameters to understand the extent of modulation of gut microbiota during antibiotic treatment (3, 35). In the current study, the Shannon diversity index at the phylum level showed a decrease in the diversity of gut microbiota in all three groups of antibiotics treated mice compared to the control group of mice. Vancomycin and neomycin treatment caused a similar extent of reduction in the diversity of gut microbiota. However, among the three antibiotic-treated groups, AVNM treatment caused a maximal reduction in gut microbial diversity (Fig 2A). The genus-level analysis of the data further validated the phylum level observations (Fig 2A).

**Fig 2.**
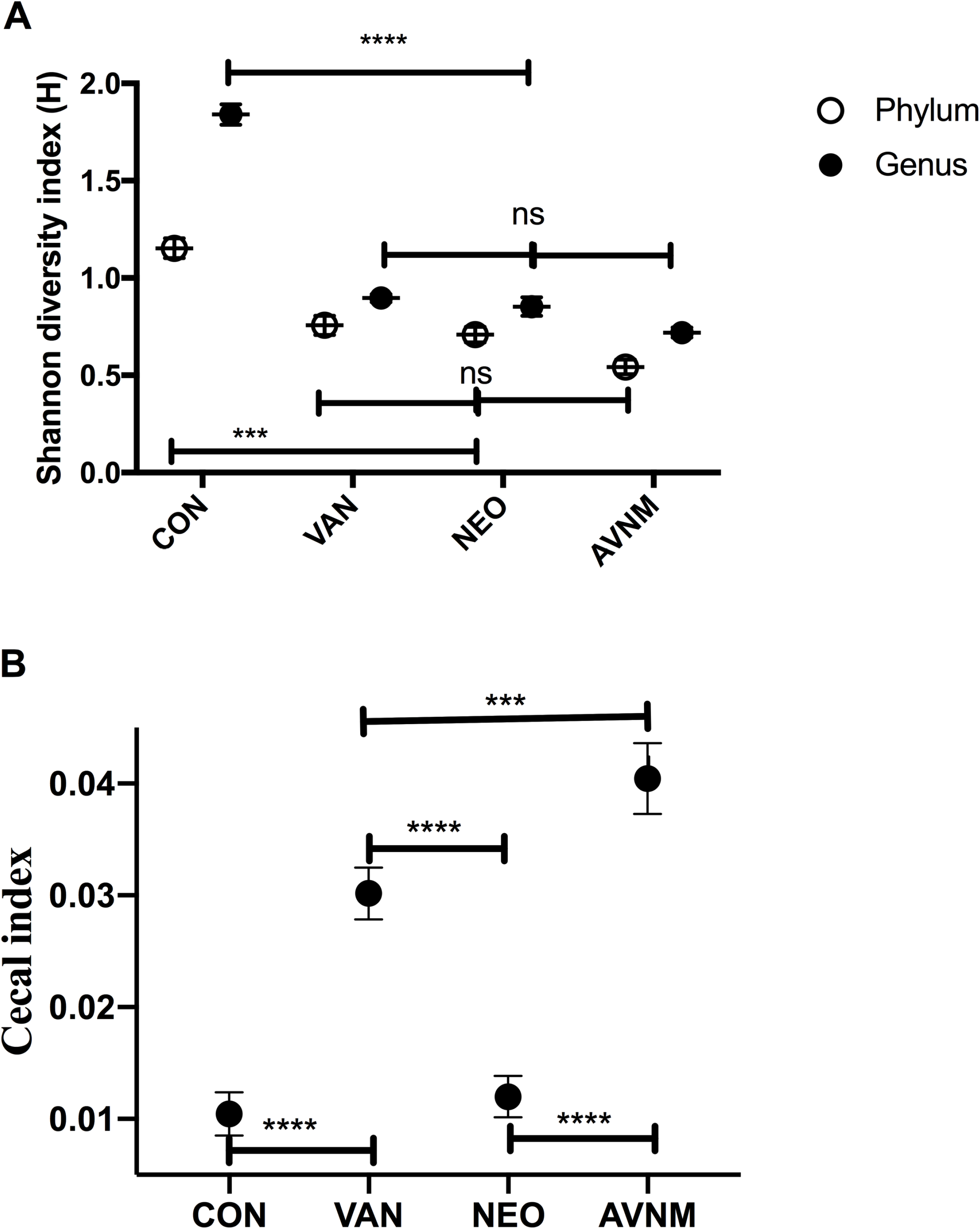
Gut microbial diversity and cecal index profile of control and different antibiotics treated mice. Shannon diversity Index for A. phylum and genus level of the cecal sample in the control and treated mice (vancomycin, neomycin, AVNM) following 7 days of treatments. B. The cecal index (ratio of the weight of the cecal content to the bodyweight) of control and antibiotic-treated mice. The statistical significance of diversity was calculated by two-way ANOVA. (‘****’ corresponds to P ≤ 0.0001,‘***’ corresponds to P ≤ 0.001, ‘**’ corresponds to P ≤ 0.01, ‘*’ corresponds to P ≤ 0.05 level of significance). Error bars shown are one standard deviation of the mean value and determined from the average values of three biological replicates.

### Alteration in the cecal index and bodyweight of mice during treatment with antibiotics

Alteration in the cecal size is usually a good indication of the variation of bacterial abundance in the cecum of mice (36). In this study, the cecal index of the antibiotic-treated mice varied significantly compared to the control group of mice (Fig 2B). It increased dramatically following vancomycin and AVNM treatment. AVNM treated group had the highest weight of cecal content among all groups of mice. However, the neomycin treated group of mice showed no changes in the cecal weight compared to the control group of mice.

We have measured the bodyweight of control and antibiotic treated mice from day zero to day seven of the experiment but could not find any significant difference between starting (day zero) and ending points (day seven) of the experiment (Table 2). Fluid consumption (ml/day) of AVNM treated mice also did not change significantly between day zero and day seven of treatment (Table 3).

**Table 2:** Body weight of mice during antibiotic treatment.

|  | Control | Vancomycin | Neomycin | AVNM |
| --- | --- | --- | --- | --- |
| 0 | 26.43±0.3 | 25.36±0.38 | 27.4±0.9 | 26.77±0.55 |
| 1 | 26.4±0.36 | 25.17±0.28 | 27.28±0.75 | 26.2±0.46 |
| 2 | 26.36±0.32 | 25±0.62 | 26.81±0.16 | 25.8±0.75 |
| 3 | 26.5±0.4 | 24.85±0.58 | 27.1±0.75 | 25±0.3 |
| 4 | 26.58±0.23 | 24.7±0.5 | 27.26±0.54 | 25.9±0.37 |
| 5 | 26.37±0.29 | 25.13±0.35 | 27.2±0.25 | 26.34±0.43 |
| 6 | 26.42±0.18 | 25.41±0.39 | 27.5±0.3 | 26.61±0.58 |
| 7 | 26.51±0.25 | 26±0.7 | 28±0.5 | 26.58±0.67 |

Body weight of control and antibiotic-treated mice (Vancomycin, Neomycin, AVNM) from day zero to day seven were shown. Means of body weight (g) of mice on different days with their respective standard deviations ( ± SD) were represented.

**Table 3:** Daily Fluid consumption (ml/day) of mice during AVNM treatment

| Days | Control | AVNM |
| --- | --- | --- |
| 0 | 5.5±0.45 | 5.8±0.5 |
| 1 | 5.7±0.35 | 5.21±0.25 |
| 2 | 5.36±0.47 | 4.8±0.45 |
| 3 | 5.81±0.24 | 4.64±0.58 |
| 4 | 5.6±0.36 | 5±0.36 |
| 5 | 5.5±0.55 | 5.46±0.29 |
| 6 | 5.8±0.4 | 5.74±0.4 |
| 7 | 5.9±0.65 | 5.3±0.5 |

Daily fluid consumption kinetics for control and AVNM-treated mice from day zero to day seven were shown. Means of fluid consumption (ml/day) of mice with their respective standard deviations ( ± SD) were represented.

### The inflammatory response in the colon changed following antibiotics mediated microbiota perturbation

Gut microbiota composition and diversity regulated the expression of various Immune genes in the gut (37). Different immune genes were either up- or down-regulated depending on the antibiotic treatment groups.

Variation in the expression of select immune genes in the colon of mice following treatment with antibiotics was determined by the qRT-PCR (Fig 3). Vancomycin treated group of mice showed an increase in IL-10 gene expression in the colon, but no significant changes were observed in the expression of TNF-α and IFN-γ genes. While neomycin treated group of mice showed increased in both IL-10 and IFN-γ expression in the colon. AVNM treatment caused an increase in TNF-α gene expression in the colon but no significant changes were found in the expression of IL-10 and IFN-γ genes (Fig 4).

**Fig 3.**
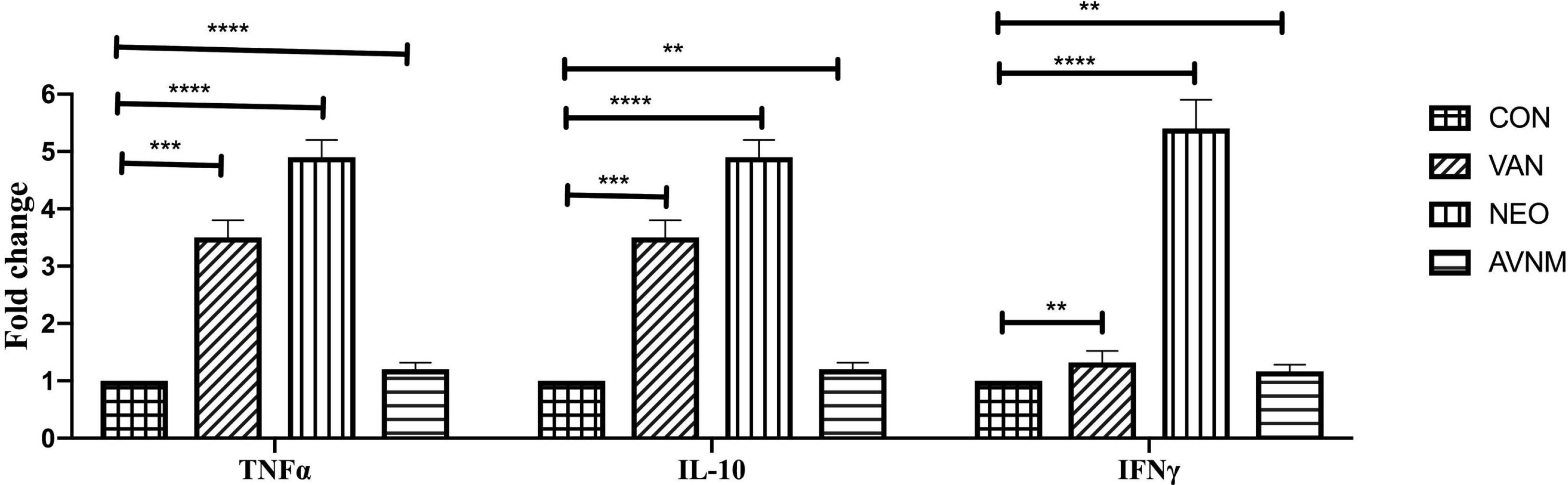
Transcriptional level expression of select inflammatory genes following treatment with different antibiotics. Fold change values of TNF-α, IFN-γ, and IL-10 are shown at mRNA level by qRT-PCR following treatment with either vancomycin (VAN), or neomycin (NEO), or AVNM for 7 days. Fold change values were calculated with respect to the untreated control expression. Control fold change value was normalized to 1. Statistical significance of the difference was calculated by two-way ANOVA (‘****’ corresponds to P ≤ 0.0001, ‘***’ corresponds to P ≤ 0.001, ‘**’ corresponds to P ≤ 0.01, ‘*’ corresponds to P ≤ 0.05 level of significance). Error bars shown are one standard deviation of the mean value and determined from the average values of four biological replicates.

**Fig 4.**
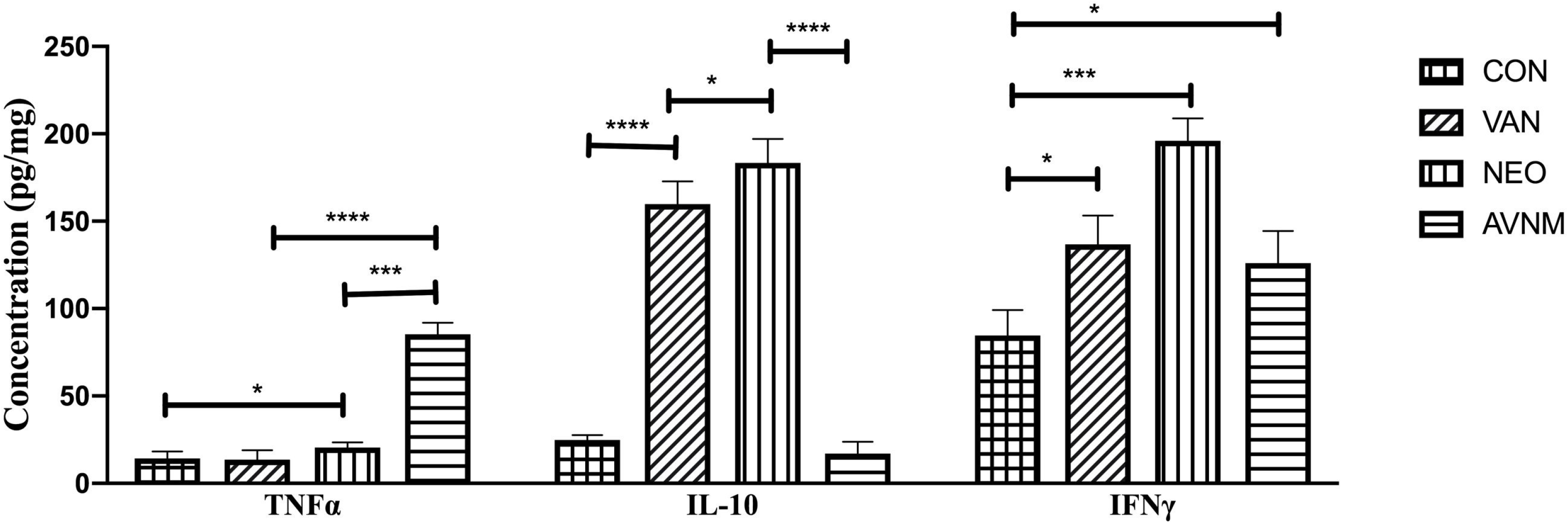
Protein level expression of select inflammatory genes in the mice gut following treatment with different antibiotics. Protein level validation for TNF-α, IFN-γ, and IL-10 abundance in the colon tissue. Statistical significance of the difference was calculated by two-way ANOVA (‘****’ corresponds to P ≤ 0.0001, ‘***’ corresponds to P ≤ 0.001, ‘**’ corresponds to P ≤ 0.01, ‘*’ corresponds to P ≤ 0.05 level of significance). Error bars shown are one standard deviation of the mean value and determined from the average values of four biological replicates.

Validation of qRT-PCR results at the protein level was done by ELISA by measuring the expression of TNF-α, IFN-γ, and IL-10 genes in the samples of colon tissue (Fig 4) and serum of the host (Fig 5). ELISA results revealed that the expression of TNF-α was the highest in the colon (Fig 4) and serum (Fig 5) of AVNM treated mice whereas the abundance of IL-10 was more in both neomycin and vancomycin treated mice (Figs 4 and 5). IFN-γ concentration was the highest in the serum sample of neomycin treated mice (Fig 5). The ELISA data for immune genes validated the qRT-PCR results.

**Fig 5.**
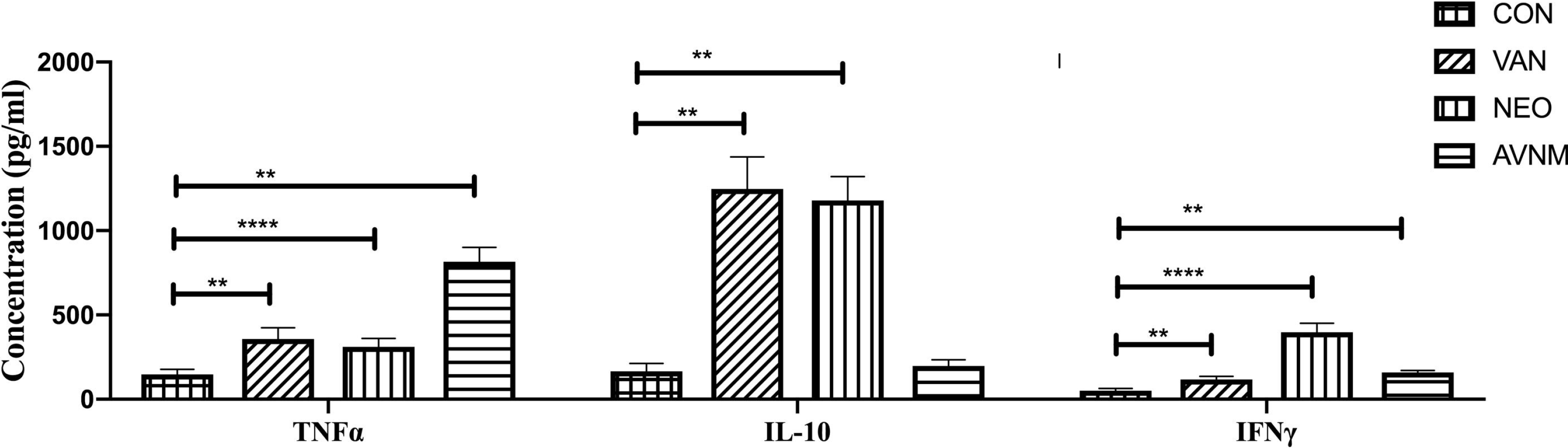
Protein level expression of select inflammatory genes in the mice serum following treatment with different antibiotics. Expression abundance of TNF-α, IFN-γ, and IL-10 in the serum of mice are shown by ELISA. Statistical significance of the difference was calculated by two-way ANOVA (‘****’ corresponds to P ≤ 0.0001, ‘***’ corresponds to P ≤ 0.001, ‘**’ corresponds to P ≤ 0.01, ‘*’ corresponds to P ≤ 0.05 level of significance). Error bars shown are one standard deviation of the mean value and determined from the average values of four biological replicates.

### Variation in SCFA abundance following treatment with antibiotics

Antibiotic treatment can drastically alter the abundance of Short-chain fatty acids (SCFAs). SCFAs are important regulators of host immune processes(9). Butyrate is mainly produced by the Firmicutes phylum while acetate and propionate are mainly produced by the Bacteroidetes phylum (9). Some earlier reports showed that *Akkeremansia muciniphila* also produced acetate and propionate in the gut (11).

We measured the concentrations of SCFAs in the host serum using NMR based metabolomics study. Results revealed that neomycin treatment caused the highest increase in propionate and acetate level with a significant decrease in butyrate level compared to control mice (Fig 6). Vancomycin treated mice showed a decrease in acetate and butyrate level compared to control mice while no significant changes were found in the propionate level. However, AVNM treated mice showed the most significant decrease in all three SCFAs, such as acetate, propionate, and butyrate compared to control and the other two antibiotics treated groups of mice (Fig 6). We also measured the abundance of acetate in the serum of both antibiotic-treated and control groups of mice by using an acetate colorimetric assay kit (EOAC-100, San Francisco, USA). The results showed that the concentration of acetate through the colorimetric detection kit method for different groups of mice, like control (51.2±4 μm), vancomycin (42±6 μm), neomycin(60±6.3 μm), and AVNM (20±1.4μm)showed nearly similar trends with NMR data.

**Fig 6.**
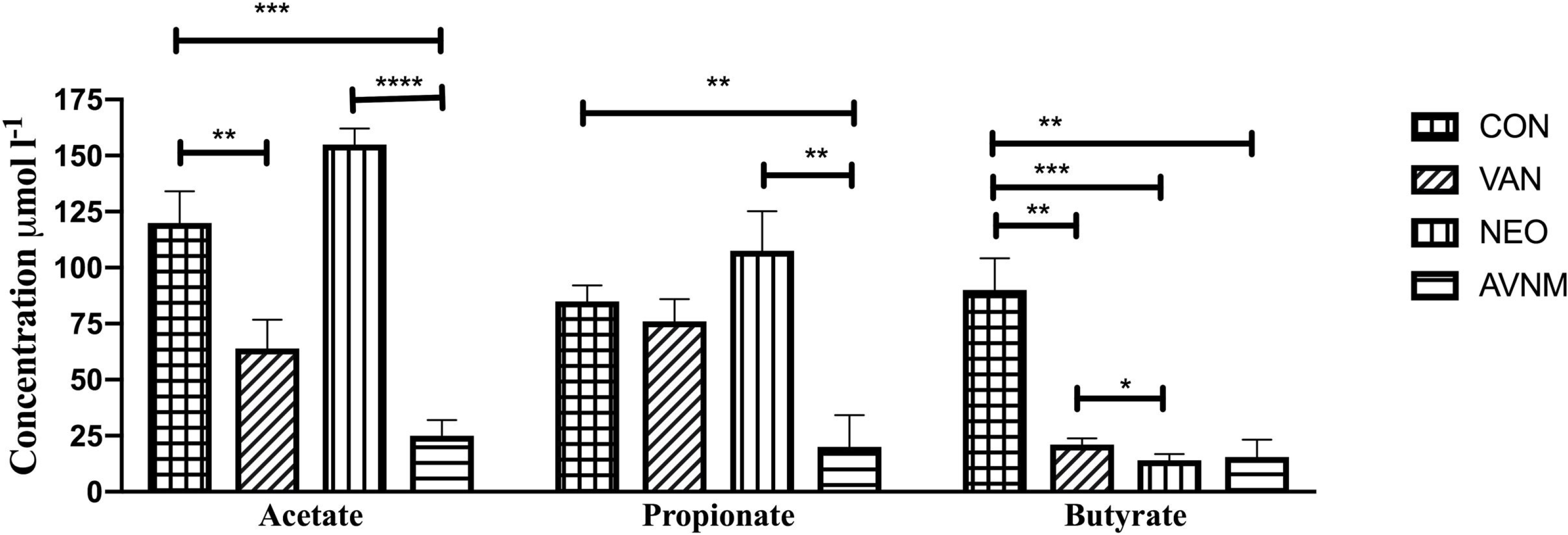
The concentration of three key short-chain fatty acids in the serum of control and different antibiotics treated mice. The concentrations of major short-chain fatty acids are shown for A. acetate, B. butyrate, and C. propionate in the serum of untreated control (con), and in the mice following 7-days of treatment with vancomycin (VAN), or neomycin (NEO), or AVNM from^1^H-NMR studies. Statistical significance of the comparison was calculated by two-way ANOVA (‘***’ corresponds to P ≤ 0.001, ‘**’ corresponds to P ≤ 0.01, ‘*’ corresponds to P ≤ 0.05 level of significance). Error bars shown are one standard deviation of the mean value and determined from the average values of three biological replicates.

## Discussion

The dysbiosis pattern of gut microbiota varied significantly among the different groups of antibiotic-treated mice. Treatment, with either individual or cocktail of antibiotics, yielded a significant yet differential in the diversity of gut microbiota. We noticed a significant increase of a particular single but distinct phylum following each type of antibiotic treatment. For example, Vancomycin treatment caused a significant increase in Verrucomicrobia phylum. The treatment with neomycin yielded an increase in the Bacteroidetes phylum, while the treatment with AVNM led to an increase in Proteobacteria phylum with lowest diversity of microbiome in the gut when compared to the other two antibiotics treatment conditions.

AVNM treatment was found to be the most effective one to reduce the diversity of gut microbiota, therefore this cocktail was taken as one of the standard gut microbial depletion agents (38–41). Contrary to the literature (34, 42), the current study showed that AVNM treatment caused extensive depletion of gut microbiota but not completely or not to the extent to make a pseudo-gnotobiotic mice. Antibiotic treatment also promotes outgrowth of resistant bacteria that make it different from germ-free mice which are free of all microorganisms (33, 43). In this study, we found that AVNM treatment caused a significant increase in the abundance of Proteobacteria phylum which replaced the other major phyla of gut microbes like Firmicutes and Bacteroidetes.

A strong correlation was found between the altered abundance of the specific gut microbes and the expressions of various immune genes in the colon of mice. Increased Akkermansia, Bacteroidetes, Escherichia-Shigella like genera, and decreased Clostridia like genus following antibiotics treatment caused significant modulation in the expression of various immune genes in the colon. Altered levels of Firmicutes and Bacteroidetes in the gut also differentially regulated the serum SCFA concentration in each antibiotic-treated group. Following vancomycin treatment, an increased abundance of Akkermansia and Lactobacillus genera caused the increased expression of the anti-inflammatory IL-10 gene in the colon of mice. Whereas no significant changes were found in the expression of pro-inflammatory genes like TNF-α and IFN-γ. Previous studies showed that the increased abundance of *Akkermansia muciniphila* induced elevated expression of anti-inflammatory cytokine genes in the gut (24, 25, 44). *A. muciniphilla* produces SCFA like acetate and propionate (9). In this study, the serum of vancomycin treated mice showed comparatively lower concentration of butyrate than propionate and acetate which could be a result of decreased Firmicutes (specifically intestinimonas) and increased *A.muciniphila* bacteria in the gut post vancomycin treatment (24, 25).

Following neomycin treatment, a significant increase in the Bacteroides genus of Bacteroidetes phylum caused an increase in the expression of both IFN-γ and IL-10 genes. It was already reported that the increased abundance of *Bacteroides fragilis* caused alteration in the expression of various immune genes of the gut tissue (45–47). Some selected gram-negative bacteria in the gut stimulated the production of IL-10 cytokine (46). It is commonly known that bacteria from the Firmicutes phylum produce butyrate while the Bacteroidetes phylum produces acetate and propionate from dietary fibers (9). In this study, following neomycin treatment, a significant reduction of Firmicutes and elevation of Bacteroidetes phylum could be related to decreased butyrate with increased acetate and propionate concentration in the serum of mice. The Acetate in the host regulated different inflammatory responses of the host. It increased the expression of the IFN-γ gene by normalizing the IFN-γ promoter, activating acetylation of histone and chromatin accessibility by acetyl-CoA synthetase (ACSS)-dependent manner (48–50). Acetate treatment also increased the IL10 level of the host, while it inhibited the LPS induced TNF-α secretion in the peripheral blood mononuclear cells (PBMCs) of mice (9, 51, 52). This showed the anti-inflammatory effect of acetate supplement on the host. In the current study, neomycin treatment caused the elevated release of acetate, which can be associated with higher IFN-γ and IL10 gene expression in the mice.

Following AVNM treatment, the dramatic increase in the Pathogenic Proteobacteria like *E.coli*, Shigella, and a decrease in the Clostridia group of bacteria caused an increase in TNF-α gene expression. Whereas no significant changes were found in the expression of IFN-γ and IL-10 genes. Previous reports showed that Firmicutes, specifically the Clostridium group present in the gut produced short-chain fatty acids and these SCFAs suppressed the LPS and pro-inflammatory cytokines (10, 11). Earlier reports showed that a considerable increase in the *Escherichia coli* like pathogenic Proteobacteria caused the higher expression of pro-inflammatory cytokine genes in the gut (53–55). In the current study, due to a significant reduction in major phyla like Firmicutes and Bacteroidetes, we observed a substantial decrease in all three SCFAs (acetate, propionate, and butyrate) level in the serum of AVNM treated mice compared to control and other antibiotic treated groups.

Bacteria of Intestinimonas genus (Firmicutes phylum), produce butyrate and Bacteroidetes produces propionate in the gut (56, 57). The production of these SCFAs in the gut suppresses the LPS and pro-inflammatory cytokines like TNF-α level and enhances the release of the anti-inflammatory cytokine like IL-10 in the colon (10, 11). In the current study, AVNM treatment caused a decrease in the concentrations of all three SCFAs which can be associated with higher TNF-α and lower IL-10 level in the colon of mice. In neomycin and vancomycin treated mice, a higher level of propionate and acetate caused more production of anti-inflammatory cytokine-like IL10 compared to AVNM treated mice.

The current study established the differential nature of gut microbial dysbiosis following treatment with either vancomycin or neomycin or AVNM in C57BL/6 male mice. The results correlated treatment of select antibiotics with gut microbial dysbiosis and metabolite and immune responses. The present study, however, did not investigate the comprehensive mechanism by which the abundance of certain groups of gut microbiota regulated the expression of different cytokines and SCFA levels of the host.

In conclusion, the current study showed different antibiotic-induced alteration patterns of gut microbiota and their association with various cytokines and SCFA levels of the host. Such association can be summed up as follows: Treatment with a) vancomycin enhanced verrucomicrobia phylum to enhance anti-inflammatory and insulin sensitivity (24, 25), b) neomycin increased Bacteroidetes phylum to promote anti-inflammatory response in the host and c) AVNM depleted most of the microbes with significant increase in pathogenic proteobacteria and beneficial verrucomicrobia phylum (24, 25). In a nutshell, the current observations are important in developing animal models for various infectious and metabolic disorder studies as well has the potential to translate clinically.

## Materials and methods

### Animals

All mice used in the present study were co-housed in polysulfone cage, and corncob was used as bedding material. Male C57BL/6 mice of 6-8 weeks of age were used in the present study. Food and water were provided *ad libitum*. Animals were co-housed in a pathogen-free environment with a 12h light-dark cycle at temperature 24 ± 3° with nearly 55% of humidity. All protocols were approved by the Institute Animal Ethics Committee constituted by CPCSEA (Reg. No.- 1643/GO/a/12/CPCSEA). All the animals were obtained from institutional Animal research and experimentation facility, School of Biological Science, NISER, Odisha, India. Animals were bred, grown, and used for the experiments in the same institutional animal house facility. For this study, the protocol number approved by NISER review committee was NISER/SBS/IAEC/AH-21. We didn’t use any anesthetic agents for the current study. Mice were euthanized by cervical dislocation method.

### Antibiotic treatment

C57BL/6 mice were divided into three groups: i) vancomycin ii) neomycin iii) AVNM treated group. Antibiotics were treated for seven consecutive days. Vancomycin treated group of mice were gavaged orally with vancomycin at a dose of 50 mg per kg of bodyweight, twice daily at a gap of 12h. Similarly, neomycin treated mice were also gavaged orally with neomycin at a dose of 50mg per kg of bodyweight twice daily. The dosages were selected as per previous reports and FDA guidelines (26, 28, 58, 59).

In AVNM treated group, AVNM cocktail (MP Biomedicals, Illkrich, France) was made by mixing four antibiotics i.e., ampicillin (1gm/lit), vancomycin (500mg/lit), neomycin (1gm/lit), and metronidazole (1gm/lit) in the drinking water. This cocktail of antibiotics was changed at a gap of every two days (at 48 h interval) and freshly prepared AVNM mixture was added in the drinking water bottle of mice. The dosage of AVNM treatment was selected as per previous reports (34, 41, 60). The AVNM mixture is a broad-spectrum antibiotic cocktail which inhibits both Gram-positive and Gram-negative bacteria of the gut. Ampicillin is one of the β-lactam antibiotics which acts against both Gram-positive and Gram-negative bacteria. Vancomycin is one of the glycopeptide antibiotics which mainly acts against Gram-positive bacteria of the intestine. Neomycin is an aminoglycoside antibiotic that has bactericidal activity against the Gram-negative bacteria and metronidazole mainly works against anaerobes. Therefore in AVNM cocktail, Ampicillin and Neomycin mainly inhibit Gram-negative bacteria of the gut while Ampicillin and Vancomycin inhibit Gram-positive bacteria which makes AVNM as an efficient antimicrobial cocktail with high depletion ability of gut bacteria (34).

Bodyweights for untreated control and all antibiotic-treated mice were recorded from the zeroth day to the seventh day of the experiment.

### Mice treatment and sample collection

Mice were separated into two different groups with Control (untreated) and Treatment (groups that were treated with antibiotics). On the 7^th^ day of the experiment, time-matched control and treated mice were euthanized by cervical dislocation. Colon tissue and cecal materials were isolated from each mouse (n=5). Tissue samples, not used immediately, were stored in RNAlater for RNA analysis until further used (61–63). Blood was collected from both control and antibiotic-treated mice for metabolomic study.

RNA extraction: RNA was extracted from the colon tissue of mice (n=5) by using the RNeasy mini kit (Cat# 74104, Qiagen, Germany) following the manufacturer’s protocol. 20-23 mg of tissue was processed using liquid nitrogen followed by homogenization in 700 μl of RLT buffer. An equal volume of 70% ethanol was added and mixed well. The solution was centrifuged at 13,000 rpm for 5 minutes at room temperature. The clear solution containing lysate was passed through the RNeasy mini column (Qiagen, Germany), which leads to the binding of RNA with the column. The column was washed using 700 μl RW1 buffer and next with 500 μl of RPE buffer. RNA was eluted using 30 μl of nuclease-free water. RNA was quantified using NanoDrop 2000 (ThermoFisher Scientific, Columbus, OH, USA).

### cDNA preparation of the extracted RNA

cDNA was synthesized from the previously extracted RNA of mice colon tissue by using Affinity Script One-Step RT-PCR Kit (Cat# 600559, Agilent, Santa Clara, CA, USA). RNA was mixed with random 9mer primers, Taq polymerase, and NT buffer. The mixture was kept at 45 °C for 30 min for the synthesis of cDNA and temperature increased to 92 °C for deactivation of the enzyme.

### Real-time PCR

(qRT-PCR) of the prepared cDNA: Real-time PCR was performed in a 96-well plate, using 25 ng cDNA as a template, 1 μM of each of forward (_F) and reverse (_R) primers for genes mentioned in Table 4, SYBR green master mix (Cat#A6002, Promega, Madison, WI, USA), and nuclease-free water. qRT-PCR was performed in Quantstudio 7 (Thermo Fisher Scientific, Columbus, OH, USA). All values were normalized with the cycle threshold (Ct) value of GAPDH (internal control) and fold change of the desired gene was calculated with respect to the control using the protocol described before(62, 63).

**Table 4:**
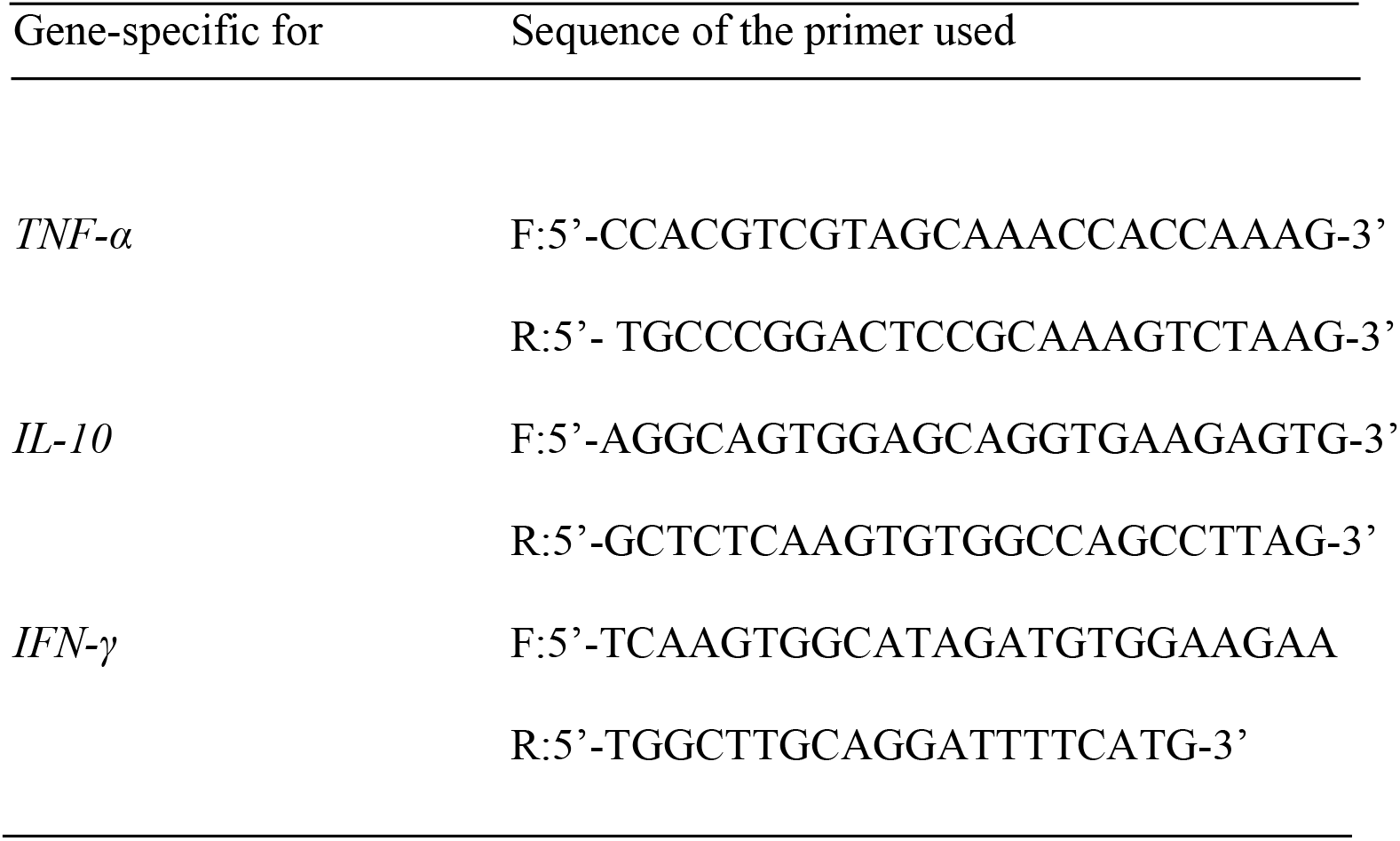
Sequences of forward (_F) and reverse (_R) primers for PCR studies to confirm the expression of various genes used in this study.

Serum collection: Mice were anesthetized (n=3) and whole blood was collected by cardiac puncture. Blood was kept on ice for 30 mins and centrifuged at 1700g for 15 min at 4°C, and serum was collected for further analysis. If required, serum was stored at −80°C until further use.

#### Cytokine Analysis at the protein level

ELISA was performed in both serum sample and colon tissue of control and each group of antibiotics treated mice. Colon tissues were collected from each antibiotic-treated and untreated (control) group of mice following seven days of treatment. After washing the colon tissues thoroughly, lysis buffer (Tris-hydrochloric acid, sodium chloride, and Triton X-100 in distilled water) containing 1X protease inhibitor cocktail (PIC) (Cat#ML051, Himedia, India) was used to churn the tissue (61). The supernatant was collected following centrifuging the churned mixture at 20,000g for 20 minutes. ELISA (BD Biosciences, San Diego, CA, USA) was performed using the manufacturer’s protocol for TNF-α (Cat#560478), IFN-γ (Cat#551866), and IL-10 (Cat#555252) expression. Protein concentration was normalized through the Bradford assay (BD Biosciences, San Diego, CA, USA). The absorbance was taken using Multiskan Go (Thermo Fisher Scientific, Columbus, OH, USA).

Calculation of Cecal index: The bodyweight of each mouse was measured and recorded. The whole cecal content was collected in a microfuge tube and weighed for each mouse. The cecal index was measured by taking the ratio of cecal content to the bodyweight of each mouse (n=5)(36).

### Genomic DNA extraction

Cecal sample was collected from untreated control and post-antibiotic-treated groups of mice (n=3) and gDNA was extracted using the phenol-chloroform method. 150-200 mg of cecal sample was used to homogenize using 1ml of 1X PBS and centrifuged at 6700g for 10 minutes. The precipitate was lysed by homogenizing it in 1ml of lysis buffer (containing Tris-HCl 0.1M, EDTA 20 mM, NaCl 100 mM, 4% SDS (at pH 8) and thereafter heating it at 80 °C for 45min. Lipid and protein parts were removed from the supernatant using an equal volume of the phenol-chloroform mixture. This process was repeated until the aqueous phase became colorless. DNA was precipitated overnight at −20 °C with 3 volumes of absolute chilled ethanol. Finally, it was washed with 500 μl of 70% chilled ethanol and briefly air-dried. The gDNA was dissolved in nuclease-free water and quantified by using NanoDrop 2000.

### 16S-rRNA sequencing (V3-V4 Metagenomics)

From cecal DNA samples, V3-V4 regions of the 16S rRNA gene were amplified. For this amplification, V3F:5’-CCTACGGGNBGCASCAG-3’ and V4R: 5’-GACTACNVGGGTATCTAATCC-3’ primer pair was used. In the Illumina Miseq platform, amplicons were sequenced using paired-end (250bpX2) with a sequencing depth of 500823.1 ± 117098 reads (mean ± SD). Base composition, quality, and GC content of the FASTQ sequence were checked. More than 90% of the sequences had Phred quality scores above 30 and GC content nearly 40-60%. Conserved regions from the paired-end reads were removed. Using the FLASH program, a consensus V3-V4 region sequence was constructed by removing unwanted sequences. Pre-processed reads from all the samples were pooled and clustered into Operational Taxonomic Units (OTUs) by using the de novo clustering method based on their sequence similarity using UCLUST program. QIIME was used for the OTU generation and taxonomic mapping(64). A representative sequence was identified for each OTU and aligned against the Greengenes core set of sequences using the PyNAST program (65, 66). The alignment of these representative sequences against reference chimeric data sets was done. The RDP classifier against the SILVA database was used for taxonomic classification to get rid of hybrid sequences.

#### Sample preparation for NMR data acquisition and metabolite analysis

Serum was obtained from the blood of antibiotic-treated and control groups of mice as described before. Samples were prepared for NMR analyses following the protocol described earlier (67). All NMR experiments were performed at 298K on a Bruker 9.4T (400 MHz) Avance-III Nanobay solution state NMR spectrometer equipped with a 5 mm broadband probe. Water suppression was done using excitation sculpting with gradients. Gradients having duration of 1 ms and strength of 14.9 G/cm was used for excitation sculpting. Offset optimization was performed using real time ‘gs’ mode for each sample. Sinc shaped pulse of 2 ms was used for selective excitation of the water resonance. 64 transients were recorded for each set of experiments with a moderate 5s relaxation delay to ensure complete water saturation. Topspin 2.1 was used to record and process the acquired spectra. Metabolite signals from NMR spectra were identified (targeted) and quantified using Chenomx NMR Suite7.6 (ChenomxInc., Edmonton, Canada). The spectra from the FID files were automatically phased and the baseline corrected and referenced to the DSS peak at 0 ppm through the Chenomx processor. Concentrations of metabolites were obtained through a profiler using Metaboanalyst using the methodology described elsewhere (65, 67–72). The profiler was used to assign and fit the metabolites peak from the Chenomx library and SCFAs such as acetate, butyrate, and propionate were quantified from the spectral intensities according to Chenomx guidelines.

Statistical Analysis: All the graphs were plotted using GraphPad Prism version 7.0. Statistical package in Prism was used for statistical analysis for the data to perform a ‘t’-test (to compare any 2 data sets) or ANOVA (to compare more than two datasets) as described in the text

## Acknowledgment

Authors like to acknowledge the support of Animal House in maintaining and assisting the experiments with the animals.

## Funding & payment

The current work (necessary resources to perform the experiment and the infra-structure for the laboratory) was supported by the parent institute National Institute of Science Education and Research. The current work was not supported through any extra-mural funding except the Ph.D. fellowship to PR by the Council of Scientific and Industrial Research (CSIR), Govt. of India.

